# Single-linkage molecular clustering of viral pathogens

**DOI:** 10.1101/2023.08.03.551813

**Authors:** Maryelba Soto Miranda, Ramiro Narváez Romo, Niema Moshiri

**Affiliations:** Department of Computer Science & Engineering, UC San Diego, La Jolla, CA, USA

**Keywords:** Molecular clustering, pairwise distance thresholds, single-linkage clustering, clusters, TN93

## Abstract

**Introduction:** Public health faces the ongoing mission of safeguarding the population’s health against various infectious diseases caused by a great number of pathogens. Epidemiology is an essential discipline in this field. With the rise of more advanced technologies, new tools are emerging to enhance the capability to intervene and control an epidemic. Among these approaches, molecular clustering comes forth as a promising option. However, appropriate genetic distance thresholds for defining clusters are poorly explored in contexts outside of Human Immunodeficiency Virus-1 (HIV-1).

**Methods:** In this work, using the well-used pairwise Tamura-Nei 93 (TN93) distance threshold of 0.015 for HIV-1 as a point of reference for molecular cluster properties of interest, we perform molecular clustering on whole genome sequence datasets from HIV-1, Severe Acute Respiratory Syndrome Coronavirus 2 (SARS-CoV-2), Zaire ebolavirus, and Mpox virus, to explore potential pairwise distances thresholds for these other viruses.

**Results:** We found the following pairwise TN93 distance thresholds as potential candidates for use in molecular clustering: 0.00016 (3 mutations) for Ebola, 0.00014 (4 mutations) for SARS-CoV-2, and 0.0000051 (1 mutation) for Mpox.

**Conclusion:** This study provides valuable information for epidemic control strategies, and public health efforts in managing infectious diseases caused by these viruses. The identified pairwise distance thresholds for molecular clustering can serve as a foundation for future research and intervention to combat epidemics effectively.

**Availability and implementation:** All relevant data and results can be found in the following repository: https://github.com/Niema-Lab/ENLACE-2023

## INTRODUCTION

Public health faces the ongoing mission of safeguarding the population’s health against infectious diseases caused by a great number of pathogens. Epidemiology, as an essential discipline in this field, is responsible for studying the distribution, risk factors, and prevention and control strategies of such diseases (CDC, 2021; Moreda, 2020). Nevertheless, epidemiologists face limited resources and active participation from the population to keep public health under control. An epidemic refers to the occurrence of a significant increase in the number of cases of a particular disease or health-related condition during a specific period, bringing several consequences within, such as increased morbidity and mortality; social, education, and economic disruptions (Oliva *et al*., 2021; Castañeda & Segura, 2023).

With the rise of more advanced technologies, new tools are emerging to enhance the capability to intervene and control an epidemic. Among these approaches, molecular clustering comes forth as a promising option. Wertheim *et al*. (2018) revealed how molecular clustering using HIV-TRACE (Pond *et al*., 2018) enabled the prediction of future transmissions of Human Immunodeficiency Virus-1 (HIV-1) through the use of a standardized threshold. However, appropriate genetic distance thresholds for defining clusters are poorly explored in contexts outside of HIV-1.

In this work, we explore whole genome datasets from HIV-1, Severe Acute Respiratory Syndrome Coronavirus 2 (SARS-CoV-2), Zaire ebolavirus, and Mpox virus to determine potential pairwise single-linkage clustering distance thresholds for the non-HIV pathogens.

## METHODS

Whole genome sequence datasets from HIV-1, Ebola, SARS-CoV-2, and Mpox were acquired from the NCBI Virus database. The following filters were used: Sequence Type: “GenBank”, and Nucleotide Completeness: “Complete”. The FASTA files downloaded for each virus contained a randomized subset of 2,000 of all records (with the exception of Ebola, which had a total of 604 sequences, and Mpox, which had a total of 1,796 sequences).

Subsequently, Multiple Sequence Alignment (MSA) was performed on each sequence dataset using ViralMSA v1.1.30 (Moshiri *et al*., 2021) using the NC_001802 reference genome for HIV-1, the NC_002549 reference genome for Ebola, the NC_045512 reference genome for SARS-CoV-2, and the NC_063383 reference genome for Mpox. Alignments were visualized using Jalview (Waterhouse *et al*., 2009).

Pairwise genetic distances under the Tamura-Nei 93 (TN93) model of nucleotide substitution (Tamura & Nei, 1993) were calculated using tn93 v1.0.11, which is a component of the HIV-TRACE software package (Pond *et al*., 2018).

Molecular clustering was performed on each viral dataset using the web application tool https://daniel-ji.github.io/MOLECULAR-DATA-VIS. Using the distance threshold of 0.015 for HIV-1 as a point of reference, we searched for distance thresholds for the other viruses that resulted in similar molecular cluster properties. To explore the space of possible distance thresholds for a virus with reference genome length *G*, we performed molecular clustering with a threshold of ^1^/_*G*_, followed by ^2^/_*G*_, followed by ^3^/_*G*_, etc., and we attempted to calibrate molecular clustering properties of interest (shown in the first column of **Table 1**) against the values obtained from the HIV-1 point of reference.

**Table 1:** Statistics of molecular clustering results for HIV-1, Ebola, SARS-CoV-2, and Mpox.

| Statistic | HIV-1<br>0.015 | Ebola<br>0.00016 | SARS-CoV-2<br>0.00014 | Mpox<br>0.0000051 |
| --- | --- | --- | --- | --- |
| Node Count | 1,101 | 489 | 1,238 | 1,244 |
| Singletons | 900 | 115 | 765 | 555 |
| Edge Count | 10,282 | 13,586 | 20,623 | 92,775 |
| Assortativity | 0.998885 | 0.884321 | 0.183067 | -0.121417 |
| Transitivity | 0.996970 | 0.961574 | 0.625468 | 0.752738 |
| Triangle Count | 168,699 | 447,472 | 428,714 | 8,972,203 |
| Cluster Count | 179 | 40 | 42 | 111 |
| Mean Cluster Size | 6.15 | 12.22 | 29.48 | 11.21 |
| Median Cluster Size | 3 | 4 | 3 | 2 |
| Mean Pairwise Distance | 0.004189 | 0.000045 | 0.000104 | 0.000004 |
| Median Pairwise Distance | 0.001123 | 0.000000 | 0.000101 | 0.000005 |
| Mean Node Degree | 18.68 | 55.57 | 33.32 | 149.16 |
| Median Node Degree | 7 | 30 | 10 | 22 |

## RESULTS

Pairwise TN93 distance distributions are shown in **Figure 1**. The HIV-1 distribution (Fig. **1A**) has peaks in the following ranges: 0.00–0.03, 0.08–0.11, 0.13–0.17, 0.18–0.21, and 0.26–0.31. The Ebola distribution (Fig. **1B**) has peaks in the following ranges: 0.000–0.005, 0.030–0.037, and 0.054–0.057. The SARS-CoV-2 distribution (Fig. **1C**) has peaks in the following ranges: 0.0000–0.0011, 0.0012–0.0021, and 0.0022–0.0035. The Mpox distribution (Fig. **1D**) has peaks in the following ranges: 0.0000–0.0001, 0.0002–0.0005, 0.0020–0.0022, 0.0027–0.0031, 0.0044–0.0047, and 0.0064–0.0067.

**Figure 1:**
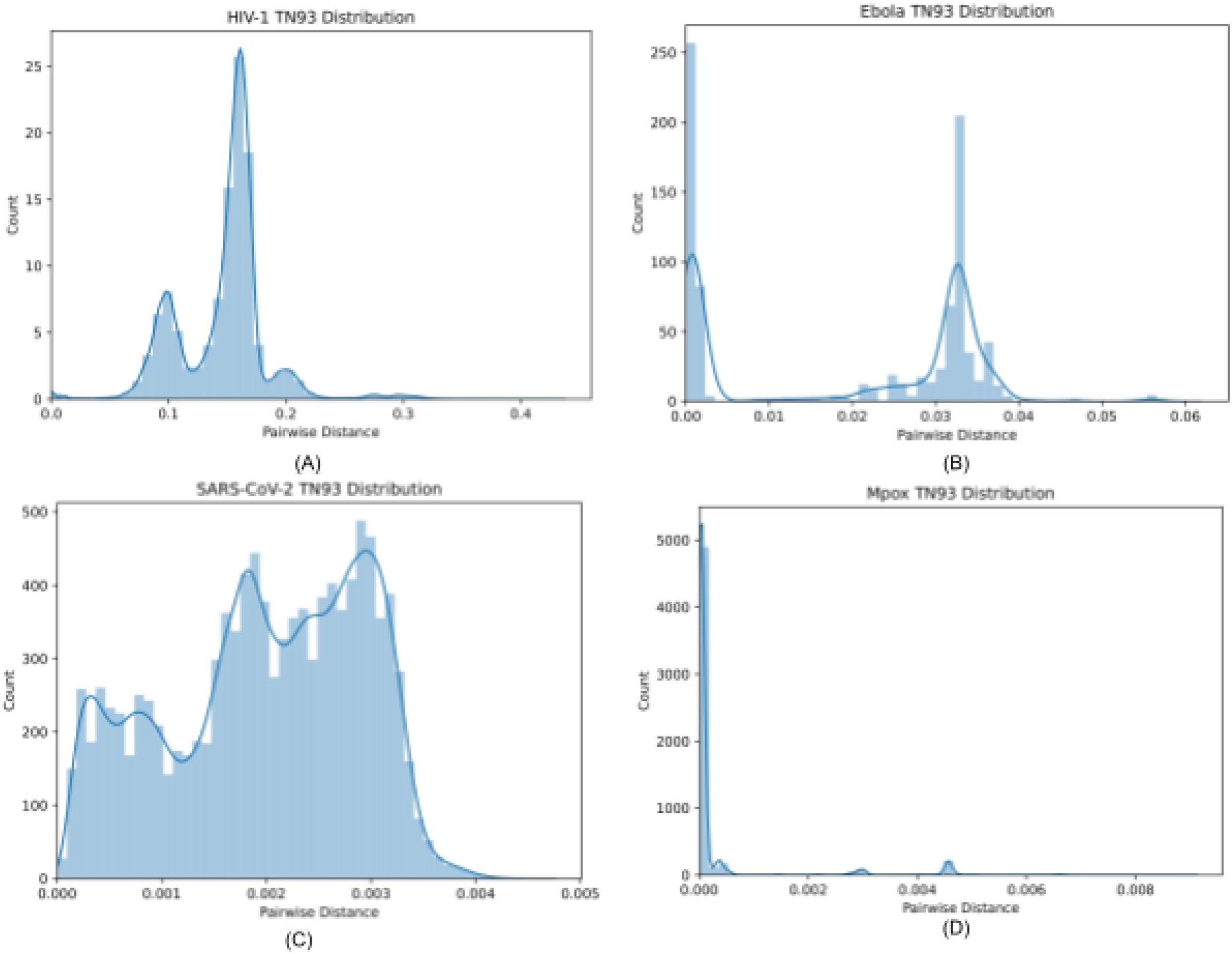
Pairwise Tamura-Nei 93 (TN93) distance distributions for (a) HIV-1, (b) Ebola, (c) SARS-CoV-2, and (d) Mpox.

After performing the calibration to the HIV-1 molecular clustering with pairwise distance threshold 0.015, we found the following potential thresholds for the other viruses: 0.00016 (3 mutations) for Ebola, 0.00014 (4 mutations) for SARS-CoV-2, and 0.0000051 (1 mutation) for Mpox. The molecular clustering properties are shown in **Table 1**, visualizations of the clusters are shown in **Figure 2**, and cluster size distributions are shown in **Figure 3**.

**Figure 2:**
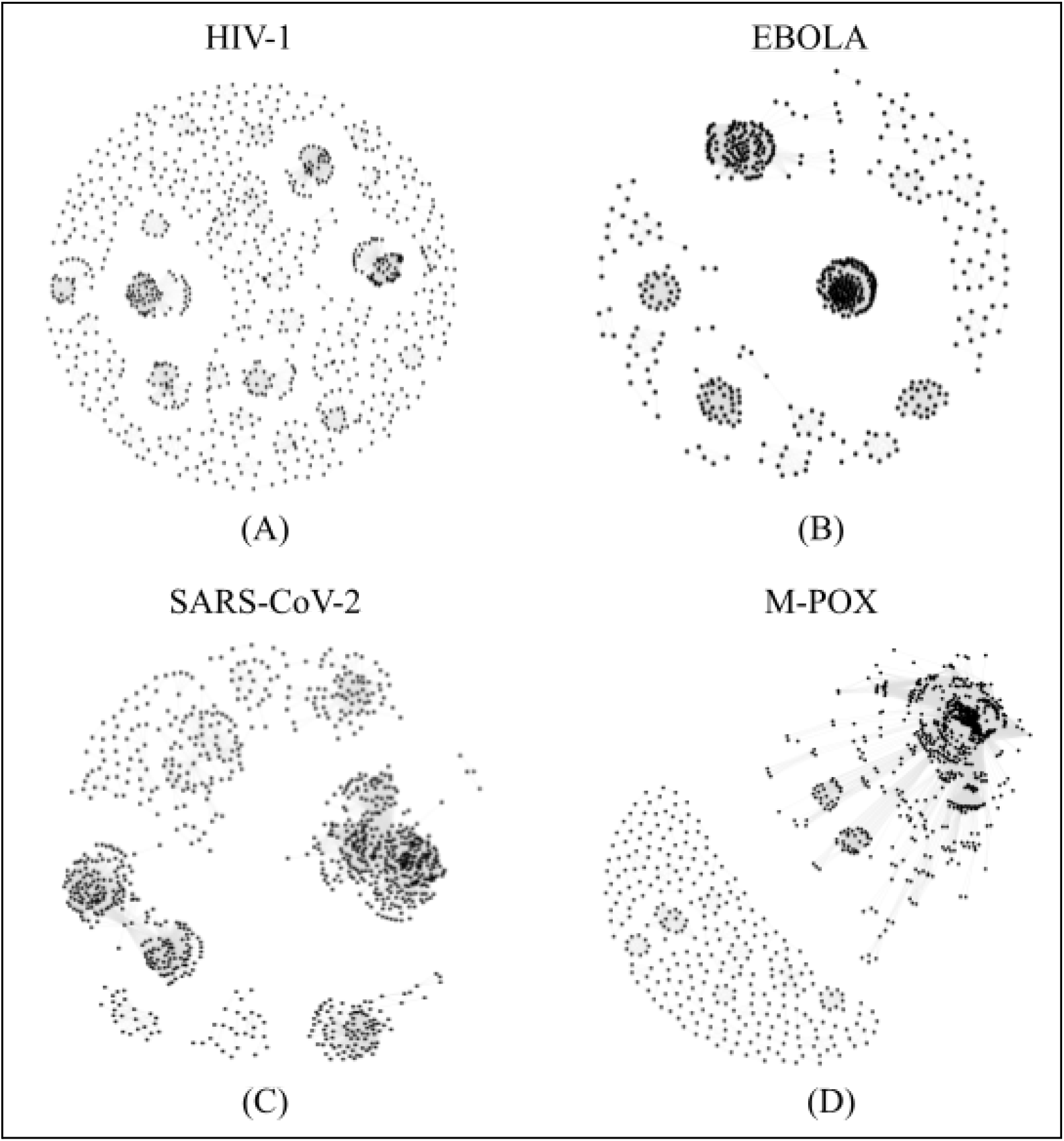
Molecular clusters of (a) HIV-1, (b) Ebola, (c) SARS-CoV-2, and (d) Mpoxn.

**Figure 3:**
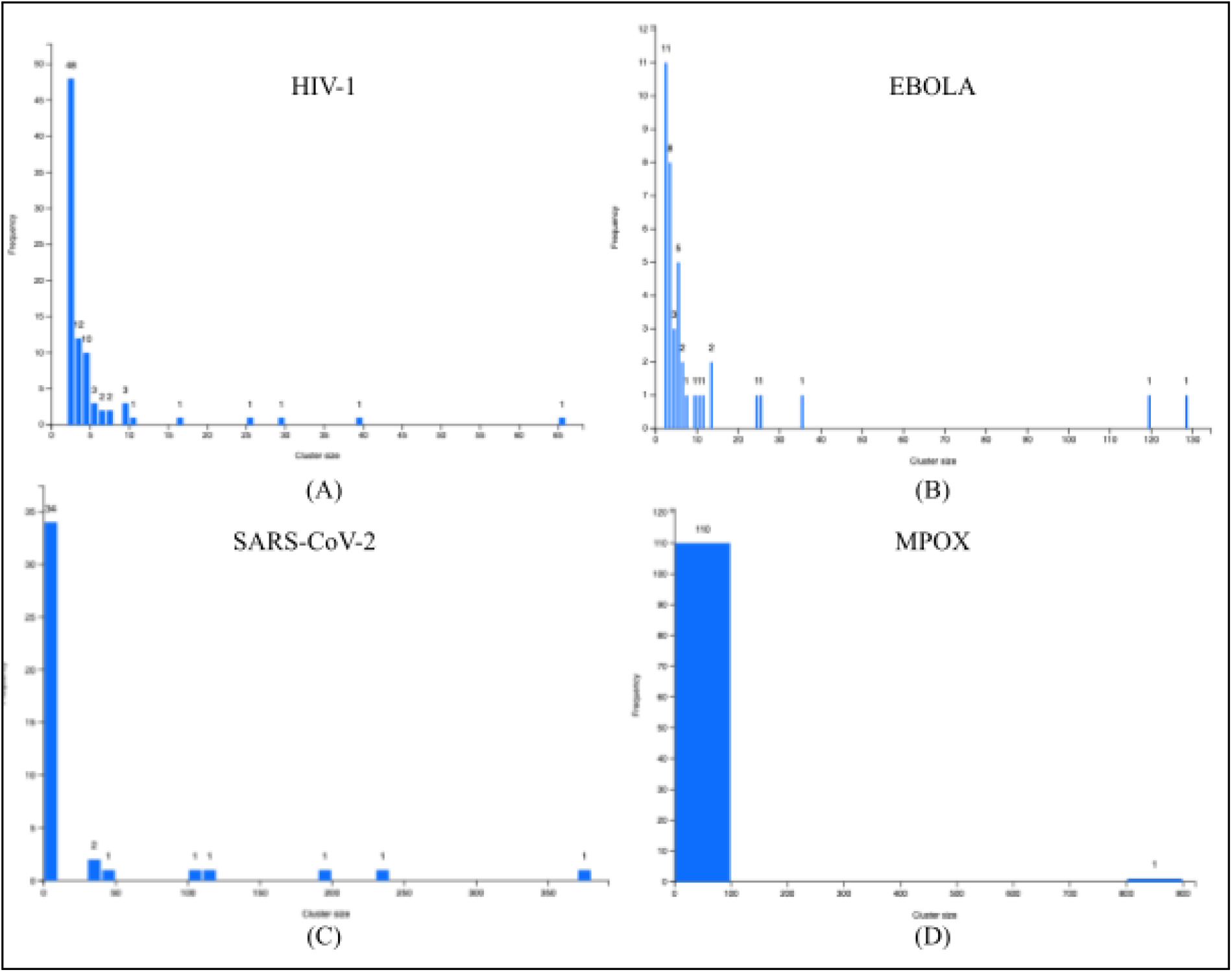
Molecular cluster size distribution for (a) HIV-1, (b) Ebola, (c) SARS-CoV-2, and (d) Mpox.

## DISCUSSION

We used HIV-1 with a molecular clustering threshold of 0.015 as a point of reference. While adjusting the threshold for other viruses, we attempted to calibrate to HIV-1 molecular clustering properties of interest. This worked reasonably well for Ebola and SARS-CoV-2, but for Mpox, even with a pairwise distance threshold corresponding to just a single mutation, we obtained a single massive cluster (Fig. **2D**). Mpox is a DNA virus, and the increased stability of DNA viruses with respect to RNA viruses may have contributed to this. For Ebola and SARS-CoV-2, we obtained similar molecular clustering thresholds (0.00016 and 0.00014, respectively), which could potentially be attributed to similar reproduction rates (R0) between these viruses (Eisenberg, 2020; Manathunga *et al*., 2023; Althaus, 2014).

Several limitations must be considered regarding this analysis. While there exist millions of genome sequences for SARS-CoV-2 and thousands for HIV-1, due to restrictions of the NCBI Virus database, this analysis only utilized a randomized subset of genomes for either virus. Further, the standard HIV-1 pairwise distance threshold of 0.015 was determined by calibrating against known partnership data (i.e., by determining a threshold such that known partners tended to be clustered together, while individuals known to not be partners tended to be in separate clusters), whereas our analysis was based solely on pairwise genetic distances and properties of molecular clusters. Additional information about known epidemiological links, such as through contact tracing, could improve future analyses.

Finally, these types of molecular clustering explorations hold immense potential in public health and epidemiological efforts (Wertheim *et al*., 2018). By continuously exploring innovative methods for monitoring viruses and diseases within populations, we can significantly enhance our ability to respond effectively to emerging threats. While there are existing tools in place for monitoring HIV, it is essential to recognize that each virus poses unique challenges. Therefore, further research and investment should be directed toward developing specialized tools and methodologies tailored to combat a broader range of viruses. Such advancements will not only aid in early detection and containment but also contribute to a more comprehensive understanding of viral dynamics and transmission patterns.

Additionally, the integration of advanced technologies, data analytics, and interdisciplinary collaborations can accelerate the progress in this field. Governments, researchers, and healthcare professionals must unite their efforts to continually refine and expand our methods for viral surveillance. By doing so, we can proactively safeguard public health, swiftly respond to outbreaks, and ultimately mitigate the impact of infectious diseases on a global scale.

## DATA AVAILABILITY

All viral genome sequences are publicly available at NCBI Virus. The complete data and analysis results can be found at: https://github.com/Niema-Lab/ENLACE-2023

## REFERENCES

Althaus, C. L. (2014). Estimating the Reproduction Number of Ebola Virus (EBOV) During the 2014 Outbreak in West Africa. PLoS Currents.

Castañeda, C., & Segura, Á. (2023). The role of epidemiology in public health: a review of its functions and applications. Revista Colombiana de Salud Pública, 23(1), 237–250. Retrieved from http://www.scielo.org.co/scielo.php?script=sci_arttext&pid=S2027-468820230001002 37

Centers for Disease Control and Prevention (CDC). (2021). Principles of Epidemiology in Public Health Practice. Retrieved from https://www.cdc.gov/csels/dsepd/ss1978/lesson1/section11.html

Eisenberg, J. (2020, February 12). How Scientists Quantify the Intensity of an Outbreak Like Coronavirus and Its Pandemic Potential. Https://Sph.Umich.Edu/Pursuit/2020posts/How-Scientists-Quantify-Outbreaks.Html.

Kosakovsky, S. L., Weaver, S., Leigh Brown, A. J., & Wertheim, J. O. (2018). HIV-TRACE (TRAnsmission Cluster Engine): a Tool for Large Scale Molecular Epidemiology of HIV-1 and Other Rapidly Evolving Pathogens. Molecular Biology and Evolution, 35(7), 1812–1819.

Manathunga, S. S., Abeyagunawardena, I. A., & Dharmaratne, S. D. (2023). A comparison of transmissibility of SARS-CoV-2 variants of concern. Virology Journal, 20(1), 1–11.

Moreda, V. P. (2020). Towards an analytical framework of the demographic and economic consequences of epidemics. Research in economic history, 3–9.

Moshiri, N. (2021). ViralMSA: massively scalable reference-guided multiple sequence alignment of viral genomes. Bioinformatics, 37(5):714–716.

Moshiri, N., Wertheim, J. O., & Mirarab, S. (2019). FAVITES: Simultaneous simulation of transmission networks, phylogenetic trees, and sequences. Bioinformatics, 35(11), 1852–1861.

Oliva, J., Delgado, C. & Laurrari, A. (2019). Guide for Severity Assessment in Influenza Epidemics. Instituto de Salud Carlos III. Retrieved from https://www.isciii.es/QueHacemos/Servicios/VigilanciaSaludPublicaRENAVE/EnfermedadesTransmisibles/Documents/GRIPE/GUIAS/Guia_Evaluacion_Gravedad_Epidemias_Gripe_28Marzo2019.pdf

Rampogu, S., Kim, Y., Kim, S. W., & Lee, K. W. (2023). An overview on monkeypox virus: Pathogenesis, transmission, host interaction and therapeutics. Frontiers in Cellular and Infection Microbiology, 13.

Tamura, K., Nei, M. (1993). Estimation of the number of nucleotide substitutions in the control region of mitochondrial DNA in humans and chimpanzees. Molecular Biology and Evolution, 10, 512–526.

Waterhouse, A. M., Procter, J. B., Martin, D. M. A., Clamp, M., & Barton, G. J. (2009). Jalview Version 2—a multiple sequence alignment editor and analysis workbench. Bioinformatics, 25(9), 1189–1191.

Wertheim, J. O., Murrell, B., Mehta, S. R., Forgione, L. A., Kosakovsky Pond, S. L., Smith, D. M., & Torian, L. V. (2018). Growth of HIV-1 Molecular Transmission Clusters in New York City. The Journal of Infectious Diseases, 218(12), 1943–1953.

